# Presence of causative mutations affecting prolificacy in the Noire du Velay and Mouton Vendéen sheep breeds

**DOI:** 10.1101/367383

**Authors:** L. Chantepie, L. Bodin, J. Sarry, F. Woloszyn, J. Ruesche, L. Drouilhet, S. Fabre

## Abstract

For many decades, prolificacy has been selected in meat sheep breeds as a polygenic trait but with limited genetic gain. However, the discovery of major genes affecting prolificacy has changed the way of selection for some ovine breeds implementing gene-assisted selection as in the French Lacaune and Grivette meat breeds, or in the Spanish Rasa Aragonesa breed. Based on statistical analysis of litter size parameters from 34 French meat sheep populations, we suspected the segregation of a mutation in a major gene affecting prolificacy in the Noire du Velay and in the Mouton Vendéen breeds exhibiting a very high variability of the litter size. After the genotyping of mutations known to be present in French sheep breeds, we discovered the segregation of the *FecL^L^* mutation at the *B4GALNT2* locus and the *FecX^Gr^* mutation at the *BMP15* locus in Noire du Velay and Mouton Vendéen, respectively. The frequency of ewes carrying *FecL^L^* in the Noire du Velay population was estimated at 21.2% and the Mouton Vendéen ewes carrying *FecX^Gr^* at 10.3%. The estimated mutated allele effect of *FecL^L^* and *FecX^Gr^*on litter size at +0.4 and +0.3 lamb per lambing in Noire du Velay and Mouton Vendéen, respectively. Due to the fairly high frequency and the rather strong effect of the *FecL^L^* and *FecX^Gr^* prolific alleles, specific management programmes including genotyping should be implemented for a breeding objective of prolificacy adapted to each of these breeds.

---

In ovine breeds raised for meat purposes, numerical productivity represents an important technical and economic lever. The objective is to reach an optimum for the economic profitability of breeding. Improvement of this numerical productivity is achieved by increasing the number of lambs born per ewe at each lambing, i.e. the prolificacy, associated with the improvement of lamb viability as well as the maternal quality. This leads to increased post-natal survival and growth rate of the lambs. For decades, genetic selection efforts have been made particularly on improving prolificacy of sheep breeds. However, prolificacy is a weakly heritable polygenic trait (h^2^= 0.05 –0.2) (see the review by Bradford (Bradford, 1985)), allowing limited genetic gain. Nevertheless in some breeds, a very large effect on ovulation rate (OR) and litter size (LS) due to single mutation in fecundity major genes (called *Fec* genes, reviewed in (Fabre *et al*., 2006)) has been demonstrated. The first evidence of the segregation of a prolificacy major gene was established in the early 1980’s in Australian Booroola Merino. This was implicated by the observation of a large variability of LS and OR in this population and the presence of extremely prolific ewes in this low prolific breed (Piper and Bindon, 1982; Davis *et al*., 1982). The causal mutation named *FecB^B^* was discovered 20 years later in the *BMPR1B* gene (Bone Morphogenetic Protein Receptor 1B) on the ovine chromosome 6 by several independent research groups (Wilson *et al*., 2001; Mulsant *et al*., 2001; Souza *et al*., 2001). This mutation was thereafter introgressed in several ovine breeds around the world for research purposes or to improve their prolificacy although these latter programmes resulted in mixed outcomes (Walkden-Brown *et al*., 2009).

Up to now, many mutations were discovered worldwide in three other major genes namely *BMP15* (known as *FecX* (Galloway *et al*., 2000)), *GDF9* (known as *FecG* (Hanrahan *et al*., 2004)) and *B4GALNT2* (known as *FecL* (Drouilhet *et al*., 2013)). In France particularly, two genetic programmes were implemented to discover and to manage mutations with major effect in order to improve the prolificacy of commercial sheep populations (Mulsant *et al*., 2003; Bodin *et al*., 2011; Martin *et al*., 2014). The introgression of the Booroola *FecB^B^* mutation was started in Mérinos d’Arles in the 1980’s. Experimental testing by the French agricultural institute INRA has estimated the effect of the mutated allele on prolificacy at one extra lamb per lambing (Teyssier *et al*., 1997). A controlled diffusion of genotyped animals in commercial flocks is now implemented in the Mérinos d’Arles population (Teyssier *et al*., 2009). In the Lacaune breed, two different mutations affecting prolificacy were discovered in the selection nucleus of the OVI-TEST cooperative, *FecX^L^* in the *BMP15* gene on the chromosome X (Bodin *et al*., 2007), and *FecL^L^* at the *B4GALNT2* locus on the chromosome 11 (Drouilhet *et al*., 2009, 2013). As soon as 2005, it was decided to eradicate the *FecX^L^* mutation inducing sterility at the homozygous state and to manage the *FecL^L^* mutation which increases LS by +0.5 lamb per lambing. The selection objective is to achieve 50% of heterozygous *FecL^L^* carrier ewes in the Lacaune OVI-TEST selection nucleus flocks (Martin *et al*., 2014; Raoul *et al*., 2017).

Beyond these genetic programmes, research of putative mutations affecting LS was undertaken in several French and foreign sheep populations, leading to the discovery of three original causal mutations affecting the *BMP15* gene in the French Grivette, the Polish Olkuska and the Tunisian Barbarine breeds (Demars *et al*., 2013; Lassoued *et al*., 2017). In contrast with the seven other known mutations in the *BMP15* gene affecting prolificacy, the homozygous Grivette and Olkuska carrier ewes are not sterile but hyper-prolific (Demars *et al*., 2013).

In the present paper, we present an analysis of LS data from 34 French and one Spanish meat sheep breeds highlighting the suspicion of a mutation in a major gene in two of them, the Noire du Velay and the Mouton Vendéen breeds. Through molecular genotyping we evidenced the segregation of mutations already known to control OR and LS in ovine breeds. Moreover, we give an early analysis of frequency and effects of these two mutations in commercial populations.

## Material and methods

### Data and statistical analysis

#### Relationship between mean and variance of LS

Data come from the OVALL French national database for meat sheep genetic evaluation and research managed by the Institut de l’Elevage (French Livestock Institute) and the Centre de Traitement de l’Information Génétique (Genetic Information Processing Center, Jouy-en-Josas, France) gathering about 12 million lambings from 1986 to 2016. We have extracted the lambing career of purebred females alive in 2005 from 34 different breeds, representing 2 353 324 natural LS obtained without hormonal synchronisation treatment of oestrus. Moreover, we have added LS data from the Spanish database for genetic evaluation of Rasa Aragonesa – UPRA-Grupo Pastores (Fathallah *et al*., 2016). Basic statistical analysis (mean and variance of the observed LS) were used to characterize each breed as well as to select the animals entering in the genotyping programme.

#### Expected frequencies of LS and variance – expected career

A subsample of the 25 most numerically important French breeds gathering 88 428 ewes with at least 5 LS records each was considered to estimate the parameters of the LS distribution. As in Bodin and Elsen (Bodin *et al*., 1989), the second order regression coefficients of each LS frequency on the mean prolificacy of the breed were estimated on the subsample of all ewes with 5 records each, excluding the Noire du Velay and Mouton Vendéen breeds as well as those known to carry a major gene for OR. These coefficients (α_i_, β1_i_, β2_i_) permitted estimation of the expected frequencies of each LS_i_ for a population of a given prolificacy (prol): LS_i_ = α_i_ + β1_i_ prol + β2_i_ prol² and consequently the expected variance which could be compared to the observed frequencies and variance. They were applied to a sample including 3 breeds without obvious major genes (Rava, Rouge de l’Ouest, Charollais), 2 breeds known to carry major genes (Lacaune and Grivette) and the two “suspected breeds” of the present study (Noire du Velay and Mouton Vendéen). According to the threshold model of LS (Gianola, 1982), and using the expected frequencies of these 7 populations, it was also possible to simulate lifetime LS data of females with 5 records each. Thus, 200 000 careers were simulated with a repeatability on the underlying variate equal to 0.20 (i.e. ∼0.15 on the observed scale). These simulations provided the expected percentage of animals with 5 lambings which exceed a given mean prolificacy (i.e. 3.0). As before, this gives the expected value in the absence of a major gene in the population was compared to the observed value.

#### Genetic parameter analyses

The test of deviation from the Hardy Weinberg equilibrium was performed using a Pearson chi square test, while the association between genotypes and prolificacy groups (defined bellow) was analysed using an exact Fisher test which can take into account the very small sample size in some categories of the contingency tables. Both tests were performed using specific functions of the R package (R Development Core Team, 2008).

The heritability 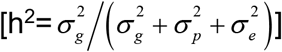 and the repeatability 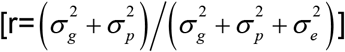 were calculated through the estimation of the additive genetic variance,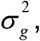 the permanent environmental variance 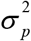 and the residual variance 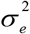. These variances were estimated by a linear mixed model run with the ASReml software (Gilmour *et al*., 2014). This model included the flock, the year, the season of lambing, the age and the genotype as fixed effects as well as random effects which were a permanent environmental effect, an animal effect whose terms were linked by a pedigree, and a residual effect. The estimation of the genotype effects were obtained from the same model by the predicted values of the genotype fixed effect.

### Animals

#### Initial sampling of Noire du Velay and Mouton Vendéen ewes based on extreme LS

The first suspicions led to selecting very small samples of relevant animals which were genotyped for the 3 known mutations naturally segregating in the French populations (*FecL^L^, FecX^L^* in Lacaune and *FecX^GR^* in Grivette). The lists of extreme highly and lowly prolific animals regarding their natural mean LS (without hormonal treatments) over at least 3 lambings were first extracted from the OVALL national database to establish the prolificacy groups. Flocks with at least 5 extreme ewes still alive at that time were selected and blood samples were collected. For the Noire du Velay breed, the final list gathered 56 females in 8 different flocks, 35 high-prolific ewes (LS mean ≥ 2.0) and 21 low-prolific ewes used as control (LS mean ≤ 1.6). In the Mouton Vendéen breed, there were 114 samples from adult ewes with 87 high-prolific (LS mean ≥ 2.20) and 27 low-prolific (LS mean ≤ 1.20).

#### Samples for studies of the frequency and the effects of the encountered mutations

In order to avoid any bias due to selection, large cohorts of unselected animals were collected in both breeds. For the Noire du Velay breed, the estimation of allele frequencies was made on unselected adult ewes (n=2728) collected in 22 different flock. After genotyping, the allele frequencies were calculated on this sample. The gene effect was estimated by a linear mixed model on the whole natural LS dataset of all ewes born after year 2000. These data (111654 records from 26398 females) as well as the pedigree of the animals were extracted from the OVALL national database. Genotypes were either unknown or determined by genotyping. The model included the flock (67 levels), the year of birth (17 levels), the age at lambing (10 levels), the season of lambing (3 levels) and the genotype (4 levels: ++, L+, LL or unknown) as fixed effects, and two random effects: a permanent environmental effect and the animal additive genetic effect.

In the Mouton Vendéen breed, blood sampling of the whole cohort of replacement ewe lambs (n=1200) belonging to 19 flocks of the selection nucleus was undertaken in 2016. A few months after sampling, these ewe lambs had their first lambing allowing estimation of the gene effect at this young age. A few adult sires were also genotyped (n=6) and the production of their daughters extracted from the national database. As for the Noire du Velay breed, the gene effect was estimated by a linear mixed model on the whole natural LS dataset of all ewes born after year 2000. These data (41269 records from 14550 females) as well as the pedigree of the animals were also extracted from the OVALL national database and the same model was applied. Levels for the fixed effects were 87 for the flock and 18 for the year of birth.

#### Blood sampling and KAPA-KASP genotyping

Blood samples (5 ml per animal) were collected from jugular vein by Venoject system containing EDTA in commercial flocks and directly stored at –20°C for further use. Genotyping was obtained by a first step of KAPA Blood PCR amplification of a specific fragment encompassing the mutation position (KAPA Biosystems). Primers used for PCR amplification were designed using Primer 3 software (Table 1). A one μl sample of total blood was run for PCR with a mixture containing 5μl of KAPA Blood kit solution and 0.25μl of each specific primer at 10nM in a final volume of 20 μl. PCR amplifications were conducted on an ABI 2400 thermocycler (Applied Biosystems) with the following conditions: 5 min initial denaturation at 94 °C, 32 cycles of 30 s at the melting temperature, 30 s extension at 72 °C and 30 s at 94 °C, followed by 5 min final extension at 72 °C.

**Table 1.**
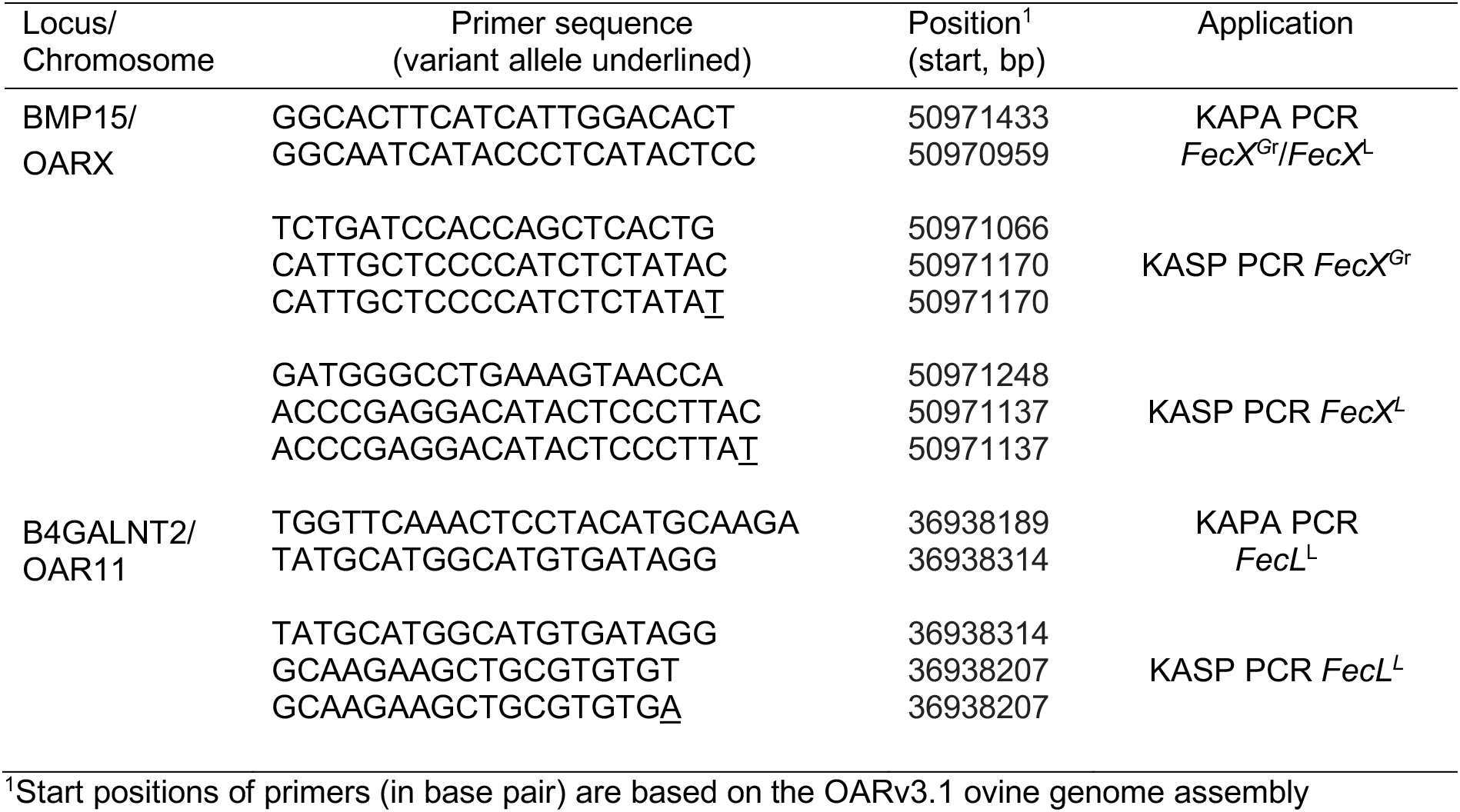
List of PCR primers used in the study.

In the second step, the specific resulting KAPA Blood PCR fragments were used as template for the genotyping of *FecL^L^* (*B4GALNT2* intron 7, OAR11:36938224T>A, NC_019468, (Drouilhet *et al*., 2013)), *FecX^L^* (*BMP15* exon 2, OARX: 50980449G>A, NC_019484, (Bodin *et al*., 2007)) or *FecX^GR^* (*BMP15* exon 2, OARX: 50980461C>T, NC_019484, (Demars *et al*., 2013)). The genotyping was done by fluorescent Kompetitive Allele Specific PCR via the KASP V4.0 2× Master mix (LGC genomics) as follow: reaction of 1.2μl of the KAPA Blood PCR product, 0.07μl primers premix and 2.5μl of the 2× KASP Master mix. The primers premix is prepared as follow: 1.2μl of each forward fluorescent allele specific primers at 100μM, 3μl of the common reverse primer at 100μM in a final volume of 10μl. Primers used for KASP PCR amplification are indicated in the Table 1. The PCR amplification condition was 15 min at 94°C for the hot-start activation, 10 cycles of 20 s at 94°C, 61- 55°C for 60 s (dropping 0.6°C per cycle), then 26 cycles of 20 s at 94°C and 60 s at 55°C. KASP genotyping was analysed by a final point read of the fluorescence on an ABI 7900HT Real-Time PCR System and using the SDS Software 2.4 (Applied Biosystems).

## Results

### Suspicion of mutation in major genes affecting prolificacy

As previously described, as long as there are less than 1% triplets in a sheep population, the distribution of LS approximately follows a binomial distribution for which the variance is directly linked to the mean (Bodin *et al*., 1989). For the breeds with a higher percent of triplets, the mean-variance relationship remains very strong. A plot of the relationship between mean and variance of LS following natural oestrus in 34 different French breeds and one Spanish breed is shown in Fig. 1. The breeds in which a mutation in a major gene is segregating were distinctly marked (Lacaune, Grivette and Rasa Aragonesa, black squares) and clearly stood out from the quadratic trendline (dashed line) which had a high r² (0.93). Excluding the three breeds with known mutations in major genes gave a quadratic trendline (plain line) with a higher r² (0.97). Based on this second trendline, the figure 1 shows that the Noire du Velay and Mouton Vendéen breeds (circles) also clearly deviated from their expected place, which could suggest the segregation of a mutation in a major gene in these two populations.

**Figure 1.**
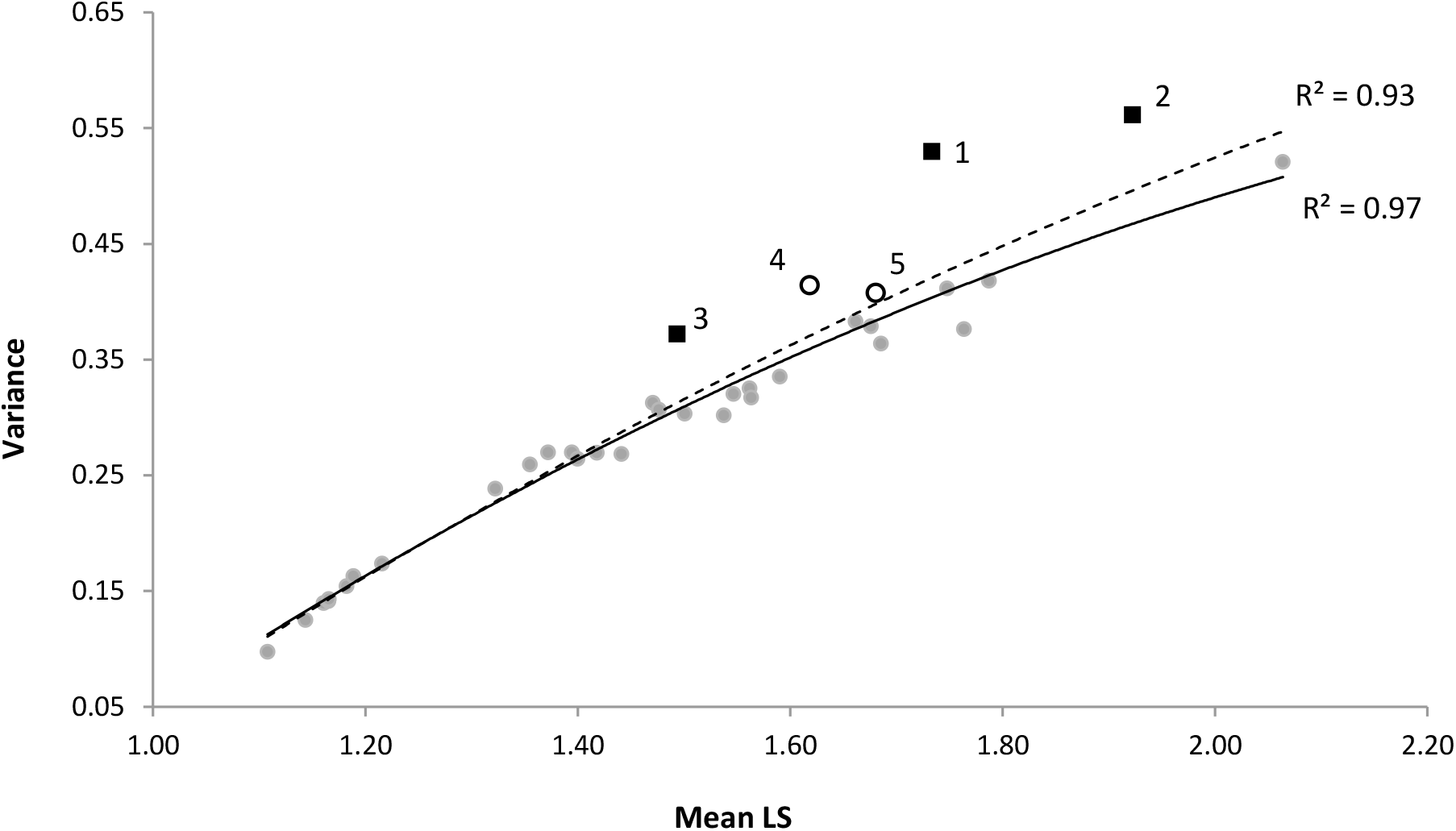
Plot of mean and variance of litter size for 35 sheep breeds. Data are from the French national OVALL database for genetic evaluation and research – Institut de l’Elevage, France and the database for genetic evaluation of Rasa Aragonesa – UPRA-Grupo Pastores, Spain (2 353 324 natural LS of purebred ewes alive in 2005). Each spot corresponds to a given population. The black squares correspond to breeds with an identified mutation in a major gene affecting prolificacy (1= Grivette, *FecX^Gr^*; 2=Lacaune, *FecX^L^* and *FecL^L^*; 3=Rasa Aragonesa, *FecX^R^*). The dashed line is the quadratic regression curve modelling all points (R^2^=0.93). The plain line is the quadratic regression modelling points without black squares (R^2^=0.97). The open circles are breeds suspected to carry a mutation in a major gene affecting prolificacy (4=Noire du Velay; 5=Mouton Vendéen).

Following this first hint, we analysed the evolution of the mean prolificacy of the 34 French breeds between 1986 and 2016. In the figure 2, we have plotted the overall annual mean LS weighted by the number of individuals in each population. We observed a regular increase of the mean LS of these breeds during the last three decades corresponding to +0.70 lamb/100 ewes/year. In contrast to the increase observed for the Lacaune, Grivette and Noire du Velay breeds, the Mouton Vendéen breed showed a slight regular decrease of its mean LS. Remarkably, the Noire du Velay breed had a strong increase of the mean LS with +1.40 lambs/100 ewes/year, twice as fast as what was observed for the overall mean and even faster than the Lacaune and Grivette breeds (respectively +0.68 and +0.75 lambs/100 ewes/year), known to carry a mutation in a major gene increasing prolificacy. Even if we could suppose that a strong improvement of the environment had occurred for improving the prolificacy of the Noire du Velay breed during the last decades, we can also speculate on the segregation of mutation in major genes influencing this trait.

**Figure 2.**
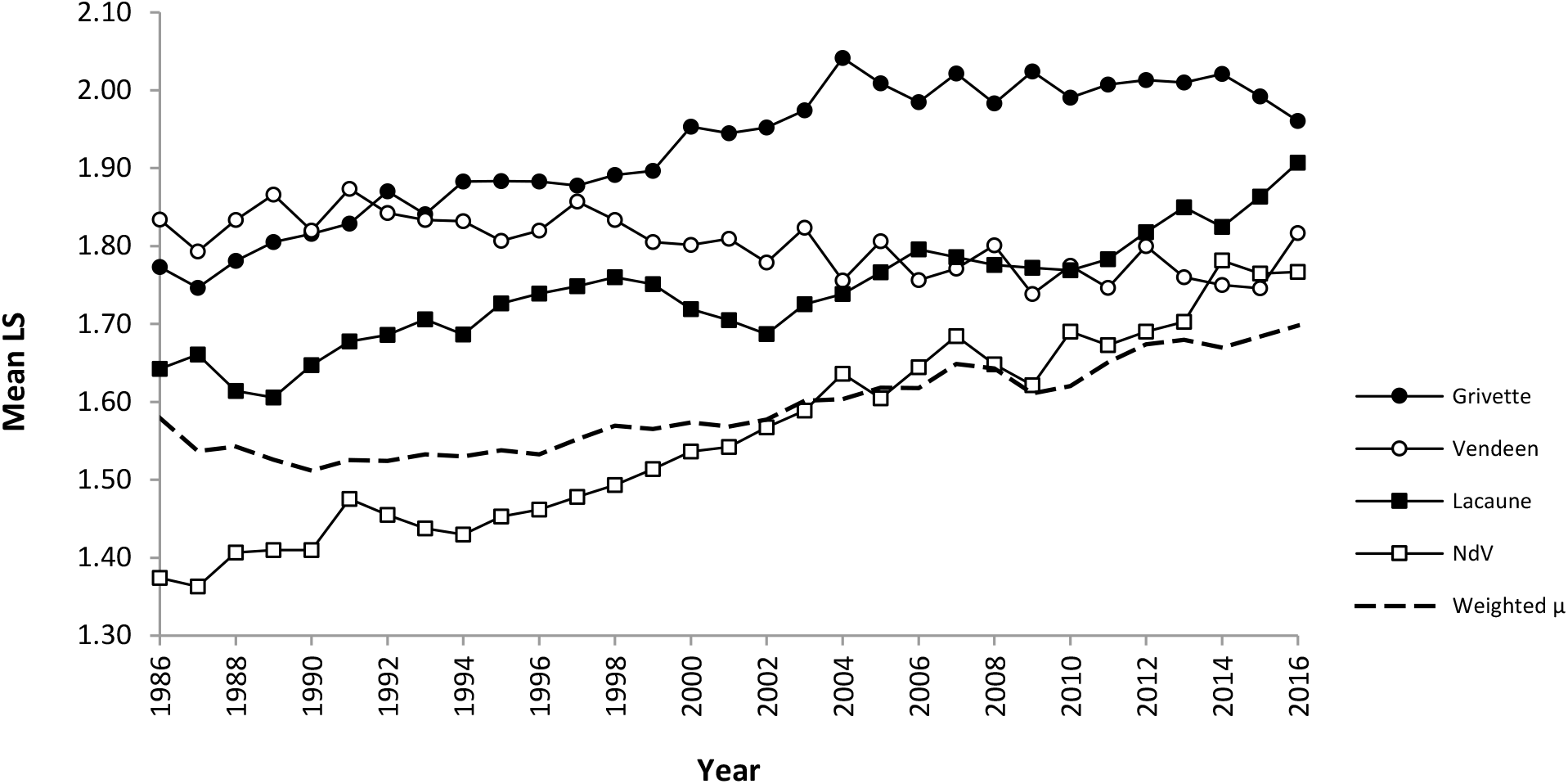
Annual evolution of mean prolificacy in French sheep breeds. Litter size (LS) data from 1986 to 2016 are from the French national OVALL database for genetic evaluation and research. The annual mean LS is plotted year by year for each breed (plain lines) and for the whole populations weighted by the number of individuals in each population (dashed line, weighted μ). NdV denotes the Noire du Velay breed.

The ratio of the observed distribution of each LS class to its expectation (provided by regression coefficients estimated on a large dataset) is given by r in Table 2. Estimations of LS frequencies were very close to the observed values for the Rava, the Rouge de l’Ouest and the Charollais breeds, as r ranged from 0.98 to 1.03. In contrast, r for the Noire du Velay and the Mouton Vendéen breeds ranged further apart from 1 (0.91 to 1.36). Similar results were obtained for the two breeds carrying a mutation in a major gene, Lacaune and Grivette (ρ ratio from 0.89 to 1.24). Consequently, ρ ratios for LS variance were remarkably close to 1 for the non-carrier breeds in contrast to those of the Lacaune and Grivette as well as the Noire du Velay and Mouton Vendéen breeds (ranging from 1.08 to 1.32). Thus, the LS distributions observed in Noire du Velay and Mouton Vendéen breeds break the rules of homogenous populations and suggested for each breed a mixture of females with different prolificacy level.

**Table 2.**
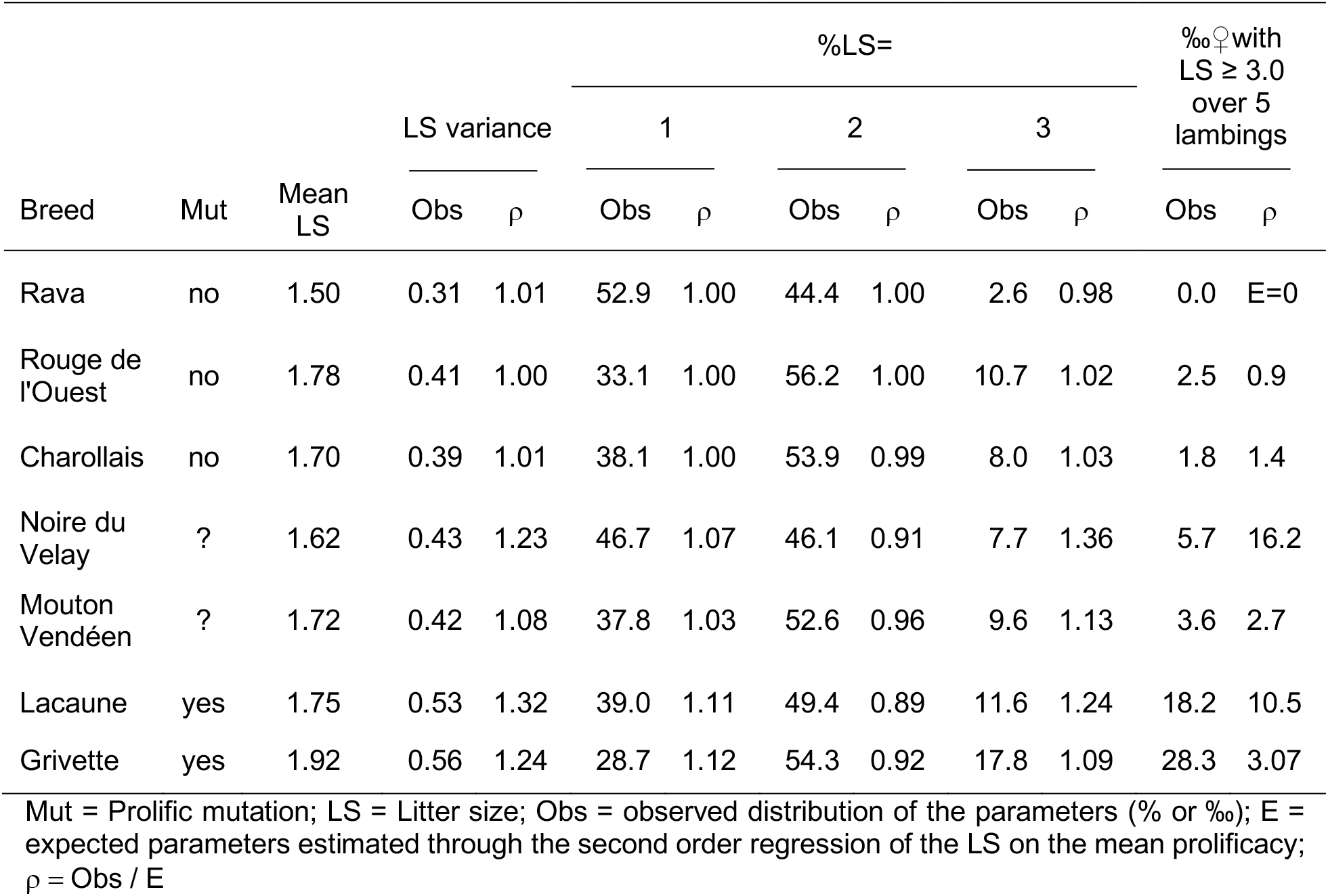
Distributions of litter size in seven different breeds carrying - or not - mutation in major gene influencing prolificacy

The final hint was the excess of highly prolific animals in these breeds (Table 2). The observed number of females having a mean prolificacy higher than 3 on five records were generally low for all the main French breeds with mean LS under 2.0. Obviously, this parameter increased regularly when the mean prolificacy of the population increased, however it was higher for the breeds known to carry or suspected to carry a mutation in a major gene reaching 28‰ in Grivette, for example (Table 2). Furthermore, r was close to 1 for the non-carrier breeds and higher for the other breeds. For the Noire du Velay breed, the number of females with a prolificacy higher than 3 was 16 times higher than expected for a comparable population without mutation in a major gene. Although this parameter was lower for the Mouton Vendéen breed, it was higher than for the non-carrier breeds.

### Genotyping of known mutations affecting prolificacy in French ovine populations: FecL^L^, FecX^L^ and FecX^Gr^

Extremely low and high-prolific ewes from the Noire du Velay (n=56) and Mouton Vendéen (n=114) populations were genotyped for the 3 known mutations naturally segregating in the French populations i.e. *FecL^L^, FecX^L^* in Lacaune and *FecX^GR^* in Grivette (Table 3). The *FecX^L^* mutation was not found but the *FecL^L^* allele was found in the Noire du Velay high-prolific group and the *FecX^Gr^* allele was found in the high-prolific group of Mouton Vendéen ewes (Table 3). All the low-prolific ewes from both breeds were wild-type at the genotyped loci. For both breeds, the exact Fisher test was highly significant (*P* < 0.001) showing a clear disequilibrium between prolificacy groups and genotypes of these mutations.

**Table 3.** FecL^L^, FecX^L^ and FecX^Gr^ genotypes of few extreme ewes for each breed and prolificacy group

| Breed | Prolific group | Locus/ Genotype |  |  |  |  |  |  |  |  |
| --- | --- | --- | --- | --- | --- | --- | --- | --- | --- | --- |
|  |  | *FecL* |  |  | *FecX* |  |  | *FecX* |  |  |
|  |  | +/+ | +/L | L/L | +/+ | +/L | L/L | +/+ | +/Gr | Gr/Gr |
| Noire du Velay | Low (n=21) | 21 | 0 | 0 | 21 | 0 | 0 | 21 | 0 | 0 |
|  | High (n=35) | 10 | 23 | 2 | 35 | 0 | 0 | 35 | 0 | 0 |
| Mouton Vendéen | Low (n=27) | 27 | 0 | 0 | 27 | 0 | 0 | 27 | 0 | 0 |
|  | High (n=87) | 87 | 0 | 0 | 87 | 0 | 0 | 56 | 29 | 2 |

### FecL^L^ and FecX^Gr^ genotype frequency and effect on prolificacy

Large cohorts of unselected animals were genotyped in order to accurately estimate the allele frequencies in the Noire du Velay and the Mouton Vendéen populations (Table 4). In Noire du Velay, the frequency of the *L* prolific allele at the *FecL^L^* locus was 0.11 with 20.1% heterozygous and 1.1% homozygous carriers. These frequencies are in Hardy Weinberg equilibrium (*P* = 0.36). Among the Mouton Vendéen replacement ewe lambs, the frequency of the *Gr* prolific allele at the *FecX^Gr^* locus was 0.05 and we observed 10.3% carrier ewes (*+/Gr* and *Gr/Gr*), only 3 animals being homozygous Gr/Gr. These frequencies were also in Hardy Weinberg equilibrium (*P* = 0.84).

**Table 4.** FecL^L^ and FecX^Gr^ genotyping in the Noire du Velay and Mouton Vendéen populations

| Genotype | Breed / Locus |  |  |  |  |  |
| --- | --- | --- | --- | --- | --- | --- |
|  | Noire du Velay / *FecL* (n=2728) |  |  | Mouton Vendéen / *FecX* (n=1200) |  |  |
|  | +/+ | +/L | L/L | +/+ | +/Gr | Gr/Gr |
| Number | 2151 | 548 | 29 | 1076 | 121 | 3 |
| Frequency ( % ) | 78.8 | 20.1 | 1.1 | 89.7 | 10.3 |  |
| SE<sup>1</sup> of frequency (%) | 0.8 | 0.8 | 0.2 | 0.9 | 0.9 |  |
| Raw mean LS<sup>2</sup> | 1.58 | 2.03 | 2.18 | 1.55 | 2.01 |  |

<sup>1</sup>SE = Standard error
<sup>2</sup>LS = Litter size

The estimated genetic effects are presented in the Table 5. Genetic parameters were very similar in both breeds. Heritability (h^2^: 0.09) and repeatability (r; 0.10 to 0.14) were low and in full agreement with the classical values of these parameters for this species (Janssens *et al*., 2004). A single copy of the *FecL^L^* in Noire du Velay increased the mean prolificacy by 0.42 lamb per lambing (*P* < 0.001). The additional increase due to a second copy of the mutation in homozygous carriers was lower (0.13; *P* = 0.096). In the Mouton Vendéen breed, the effect of a single copy of the *FecX^Gr^* allele increased the prolificacy by 0.30 lamb per lambing (*P* < 0.001), while the effect of a second copy leading to a homozygous carrier, does not further increase the prolificacy (*P* = 0.485). In both breeds, as expected, females of the unknown genotype group were slightly, although not significantly, more prolific than the corresponding females known as wild-type.

**Table 5.** Genetic parameters and allelic estimated values of litter size in Noire du Velay and Mouton Vendéen breeds

| Breed | n LS | Genetic parameters |  |  |  |  |  | Allelic estimated values |  |  |  |
| --- | --- | --- | --- | --- | --- | --- | --- | --- | --- | --- | --- |
| | | n ♀ | $\sigma^2_g$ | $\sigma^2_p$ | $\sigma^2_e$ | $h^2$ | r | zz | +/+ | +/M<sup>1</sup> | M/M |
| Noire du Velay | 111654 | 26398 | 0.036 | 0.017 | 0.334 | 0.09<br>(0.005) | 0.14<br>(0.003) | 1.60<sup>a</sup> | 1.57<sup>a</sup> | 1.99<sup>b</sup> | 2.12<sup>b</sup> |
| Mouton Vendéen | 41269 | 14550 | 0.034 | 0.005 | 0.344 | 0.09<br>(0.007) | 0.10<br>(0.005) | 1.71<sup>a</sup> | 1.68<sup>a</sup> | 1.98<sup>b</sup> | 1.99<sup>b</sup> |

LS = Litter size; $\sigma^2_g$ = Additive genetic variance; $\sigma^2_p$ = Permanent environmental variance; $\sigma^2_e$ = Residual variance; $h^2$ = Heritability (standard errors are between brackets); r = Repeatability (standard errors are between brackets); zz = Unknown genotype; ++ = Homozygous wild type, M+: heterozygous, MM: homozygous mutated type.
<sup>1</sup> M denotes the mutated prolific allele within each breed (*FecL<sup>L</sup>* or *FecX<sup>Gr</sup>*)
<sup>a,b</sup> Values within a row with different superscripts differ significantly at $P < 0.001$ .

## Discussion

Simple analyses of livestock industry data suggested the segregation of a mutation in a major gene affecting prolificacy of the Noire du Velay and Mouton Vendéen breeds. Small samples of extremely high and low prolific ewes were then genotyped for known mutations and revealed the presence of causal mutations. Among the different strands of evidence, the deviation from the relationship between mean and variance of prolificacy for breeds with a mutation in a major gene comparing to other breeds was quite remarkable. Even if these parameters were calculated without any correction for variation factors, the very high number of observations for each breed, smoothed all the effects and led to a very strong relationship among non-carrying populations. Only LS after natural oestrus were extracted from the national database as using hormonal treatments to induce ovulations modifies the mean-variance relationship (Bodin *et al*.,1989). Moreover, the within breed variability of LS variance due to polygenic effects is generally very small (Bodin *et al*., 1989; Amer and Bodin, 2006; Cottle *et al*., 2016) and could not affect the mean-variance relationship at a broad scale. Finally, the observed deviation was due to the mixture into the populations of two groups of animals widely differing in prolificacy as it had been already viewed in Lacaune (Martin *et al*., 2014) and in Rasa Aragonesa breed (Fathallah *et al*., 2016).

The procedure used in the present study is relevant only for genes having large effect (about 0.5 standard deviation of the prolificacy mean). If the presence of known mutations was not found, the process would have been followed by a GWAS to compare allele frequencies between the few selected high (cases) and low (controls) prolific ewes. The selection of two small samples of very extreme animals and their analysis considering they are two states of a qualitative trait has been already successful and allowed the discovery of two new mutations for ovulation rate (Demars *et al*., 2013). However, as shown in the Table 3, some highly prolific Noire du Velay and Mouton Vendéen ewes were non-carriers of known prolific alleles. Their extreme prolificacy could be either explained by the polygenic determinism of this trait or by the segregation of another major mutation as already described in the Lacaune population carrying both *FecX^L^* and *FecL^L^* (Bodin *et al*., 2007; Drouilhet *et al*., 2009).

The frequency of carrier ewes is much higher in the Noire du Velay breed (∼20%) than in the Mouton Vendéen breed (∼10%) which can explain that the deviation from the mean-variance relationship is also higher in this breed. However, the Hardy Weinberg equilibrium is still very well conserved. This means that in both populations prolific allele frequencies are not strongly affected by selection, particularly in the Mouton Vendéen breed as shown by the mean LS evolution during the last three decades. In both breeds it seems that carrier ewes do not produce more replacement ewe lambs in spite of their higher prolificacy and that there is no preferential culling according to the genotype. In Lacaune, Hardy Weinberg equilibrium did not hold for a long time because between 1996 and 2010 the cooperative excluded animals that were too prolific and since 2011 the cooperative’s aim has been to achieve 50% of L+ ewes (Martin *et al*., 2014).

The effects of one copy of the *FecL^L^* mutation on LS is similar in the Noire du Velay (+0.42 lamb per lambing) and in the Lacaune breed (+0.47, (Martin *et al*., 2014)), and is of the same order of magnitude as the effect of most of other known major genes for prolificacy (Bodin *et al*., 2011; Jansson, 2014). However, as it has been already noted, the effect of the *FecL^L^* mutation is much higher than the effect of the *FecX^Gr^* mutation observed in the Grivette population (+0.10 lamb per lambing, (Demars *et al*., 2013)). In Mouton Vendéen, the effect of one copy of the *FecX^Gr^* mutation (+0.30 ± 0.04 lamb per lambing) seems higher than the effect of the same mutation in the Grivette population, although in this latter population, the analysis of the allele effect was not conducted on a large sample. *FecX^Gr^* homozygous carrier ewes in the Grivette or the Mouton Vendéen population are as prolific, if not more, than the heterozygous ones (present work and (Demars *et al*., 2013)), in contrast to most mutations of the *BMP15*gene which induce sterility at the homozygous state (Bodin *et al*., 2007; Demars *et al*., 2013).

## Conclusion

Based on an analysis of a very large LS dataset from 34 French meat sheep breeds and molecular genotyping, we have highlighted and evidenced the segregation of two mutations in the *FecL* and *FecX* major genes in the Noire du Velay and the Mouton Vendéen breeds. We have determined a fairly high frequency (0.05 to 0.11) and a rather strong effect (+0.3 to +0.4 lamb/lambing) of the *FecL^L^* and *FecX^Gr^* prolific alleles. This discovery should serve as a basis for implementing specific management programmes, including genotyping of reproducers, in relation with the Noire du Velay and Mouton Vendéen selection organizations in line with their breeding objective of prolificacy.

## Acknowledgments

We thank Didier Cathalan from ROM Sélection managing the Noire du Velay population and Charline Rousseau from the Mouton Vendéen selection organization, for their precious help in the planning of blood sampling. We are grateful to the breeders who made their animals available for this study. LC was supported by a PhD grant co-funded by APIS-GENE through the Proligen project and the European Funds for Regional Development (FEDER) through the Interreg POCTEFA programme in the framework of the PIRINNOVI project (EFA103/15). Part of the Noire du Velay sampling was supported by the DEGERAM project co-funded by the FEDER Massif Central, the Régions: Aquitaine, Midi-Pyrénées, Limousin and Auvergne; and the French government.

## Declaration of interest

The authors declare that they have no competing interests

## Ethics statement

The blood sampling procedure was approved (approval number 01171.02) by the French Ministry of Teaching and Scientific Research and local ethical committee C2EA-115 (Science and Animal Health) in accordance with the European Union Directive 2010/63/EU on the protection of animals used for scientific purposes.

## Data repository resources

Raw data from the OVALL database were managed by the Institut de l’Elevage (French Livestock Institute) and the Centre de Traitement de l’Information Génétique (Genetic Information Processing Center, Jouy-en-Josas, France). Derived data supporting the findings of this study are available from the corresponding author upon reasonable request.

## References

Amer PR and Bodin L 2006. Quantitative genetic selection for twinning rate in ewes. New Zealand Society of Animal Production.

Bodin L, Di Pasquale E, Fabre S, Bontoux M, Monget P, Persani L and Mulsant P 2007. A novel mutation in the bone morphogenetic protein 15 gene causing defective protein secretion is associated with both increased ovulation rate and sterility in Lacaune sheep. Endocrinology 148, 393–400.

Bodin L, Elsen JM and Station d’amelioration genetique des animaux 1989. Variability of litter size of french sheep breeds following natural or induced ovulation. Animal Production, 535–541.

Bodin L, Raoul J, Demars J, Drouilhet L, Mulsant P, Sarry J, Tabet C, Tosser-Klopp G, Fabre S, Boscher MY and others 2011. Etat des lieux et gestion pratique des gènes d’ovulation détectés dans les races ovines françaises. In 18èmes Rencontres Recherches Ruminants., pp. 393–400. Institut de l’Elevage, Paris, France.

Bradford GE 1985. Selection for litter size. In Genetics of Reproduction in Sheep (eds. R.B. Land and D.W. Robinson), pp. 3–18. Butterworths, London.

Cottle DJ, Gilmour AR, Pabiou T, Amer PR and Fahey AG 2016. Genetic selection for increased mean and reduced variance of twinning rate in Belclare ewes. Journal of Animal Breeding and Genetics = Zeitschrift Fur Tierzuchtung Und Zuchtungsbiologie 133, 126–137.

Davis GH, Montgomery GW, Allison AJ, Kelly RW and Bray AR 1982. Segregation of a major gene influencing fecundity in progeny of Booroola sheep. New Zealand Journal of Agricultural Research 25, 525–529.

Demars J, Fabre S, Sarry J, Rossetti R, Gilbert H, Persani L, Tosser-Klopp G, Mulsant P, Nowak Z, Drobik W, Martyniuk E and Bodin L 2013. Genome-wide association studies identify two novel BMP15 mutations responsible for an atypical hyperprolificacy phenotype in sheep. PLoS Genetics 9, e1003482.

Drouilhet L, Lecerf F, Bodin L, Fabre S and Mulsant P 2009. Fine mapping of the FecL locus influencing prolificacy in Lacaune sheep. Animal Genetics 40, 804–812.

Drouilhet L, Mansanet C, Sarry J, Tabet K, Bardou P, Woloszyn F, Lluch J, Harichaux G, Viguié C, Monniaux D, Bodin L, Mulsant P and Fabre S 2013. The Highly Prolific Phenotype of Lacaune Sheep Is Associated with an Ectopic Expression of the B4GALNT2 Gene within the Ovary. PLoS Genetics 9, e1003809.

Fabre S, Pierre A, Mulsant P, Bodin L, Di Pasquale E, Persani L, Monget P and Monniaux D 2006. Regulation of ovulation rate in mammals: contribution of sheep genetic models. Reproductive Biology and Endocrinology 4, 20.

Fathallah S, Alabart JL, Bodin L, Jimenez-Hernando MA, Lahoz B, Fantova E, David I and Jurado JJ 2016. Relaciones entre los efectos del gen BMP15 y los efectos poligénicos sobre la prolificidad en la raza ovina Rasa Aragonesa. Informacion Tecnica Economica Agraria 112, 45–56.

Galloway SM, McNatty KP, Cambridge LM, Laitinen MPE, Juengel JL, Jokiranta TS, McLaren RJ, Luiro K, Dodds KG, Montgomery GW, Beattie AE, Davis GH and Ritvos O 2000. Mutations in an oocyte-derived growth factor gene (BMP15) cause increased ovulation rate and infertility in a dosage-sensitive manner. Nature Genetics 25, 279–283.

Gianola D 1982. Assortative mating and the genetic correlation. Theoretical and Applied Genetics 62, 225–231.

Gilmour AR, Gogel BJ., Cullis BR, Welham SJ, Thompson R. 2014. ASReml User Guide Release 4.1 Functional Specification, VSN International Ltd, Hemel Hempstead, United Kingdom, http://www.vsni.co.uk.

Hanrahan JP, Gregan SM, Mulsant P, Mullen M, Davis GH, Powell R and Galloway SM 2004. Mutations in the genes for oocyte-derived growth factors GDF9 and BMP15 are associated with both increased ovulation rate and sterility in Cambridge and Belclare sheep (Ovis aries). Biology of Reproduction 70, 900–909.

Janssens S, Vandepitte W and Bodin L 2004. Genetic parameters for litter size in sheep: natural versus hormone-induced oestrus. Genetics Selection Evolution 36, 543–562.

Jansson T 2014. Genes involved in ovulation rate and litter size in sheep. http://stud.epsilon.slu.se/6803. Accessed 20 May 2017.

Lassoued N, Benkhlil Z, Woloszyn F, Rejeb A, Aouina M, Rekik M, Fabre S and Bedhiaf-Romdhani S 2017. FecXBar a Novel BMP15 mutation responsible for prolificacy and female sterility in Tunisian Barbarine Sheep. BMC Genetics 18, 43.

Martin P, Raoul J and Bodin L 2014. Effects of the FecL major gene in the Lacaune meat sheep population. Genetics, selection, evolution: GSE 46, 48.

Mulsant P, Lecerf F, Fabre S, Bodin L, Thimonier J, Monget P, Lanneluc I, Monniaux D, Teyssier J and Elsen JM 2003. Prolificacy genes in sheep: the French genetic programmes. Reproduction (Cambridge, England) Supplement 61, 353–359.

Mulsant P, Lecerf F, Fabre S, Schibler L, Monget P, Lanneluc I, Pisselet C, Riquet J, Monniaux D, Callebaut I, Cribiu E, Thimonier J, Teyssier J, Bodin L, Cognié Y, Chitour N and Elsen JM 2001. Mutation in bone morphogenetic protein receptor-IB is associated with increased ovulation rate in Booroola Mérino ewes. Proceedings of the National Academy of Sciences of the United States of America 98, 5104–5109.

Piper LR and Bindon BM 1982. Genetic segregation for fecundity in Booroola Merino sheep. In Proceedings of the World Congress on Sheep and Beef Cattle Breeding (eds. R.A. Barton and W.C. Smith), pp. 394–400. Palmerston North, N.Z.

R Development Core Team 2008. R: A language and environment for statistical computing. R Foundation for Statistical Computing, Vienna, Austria. ISBN 3-900051-07-0. http://www.R-project.org.

Raoul J, Swan AA and Elsen J-M 2017. Using a very low-density SNP panel for genomic selection in a breeding program for sheep. Genetics Selection Evolution 49, 76.

Souza CJ, MacDougall C, MacDougall C, Campbell BK, McNeilly AS and Baird DT 2001. The Booroola (FecB) phenotype is associated with a mutation in the bone morphogenetic receptor type 1 B (BMPR1B) gene. The Journal of Endocrinology 169, R1–6.

Teyssier J, Bodin L, Maton C, Bouquet PM and Elsen JM 2009. Biological and economic consequences of introgression of the FecB gene into the French Mérinos d’Arles sheep. In Proceedings of Helen Newton Turner Memorial International Workshop Held Pune Maharashtra India 10–12 Novemb. 2008 (eds. S.W. Walkden-Brown, J.H.J. van der Werf, C. Nimbkar and V.S. Gupta), pp. 128–134. Australian Centre for International Agricultural Research.

Teyssier J, Elsen JM, Bodin L, Bosc P, Lefevre C and Thimonier J 1997. Interet zootechnique du gene Booroola en race Mérinos d’Arles. In 4èmes Rencontres Recherches Ruminants, pp. 223–226. Institut de l’Elevage, Paris, France.

Walkden-Brown SW, Wolfenden DH and Piper LR 2009. Use of the FecB (Booroola) gene in sheep-breeding programs. In Proceedings of Helen Newton Turner Memorial International Workshop Held Pune Maharashtra India 10–12 Novemb. 2008 (eds. S.W. Walkden-Brown, J.H.J. van der Werf, C. Nimbkar and V.S. Gupta), pp. 100–110. Australian Centre for International Agricultural Research.

Wilson T, Wu XY, Juengel JL, Ross IK, Lumsden JM, Lord EA, Dodds KG, Walling GA, McEwan JC, O’Connell AR, McNatty KP and Montgomery GW 2001. Highly prolific Booroola sheep have a mutation in the intracellular kinase domain of bone morphogenetic protein IB receptor (ALK-6) that is expressed in both oocytes and granulosa cells. Biology of Reproduction 64, 1225–1235.

